# A pipeline for interrogating and engineering single-subunit oligosaccharyltransferases

**DOI:** 10.1101/155101

**Authors:** Thapakorn Jaroentomeechai, Xiaolu Zheng, Jasmine Hershewe, Jessica C. Stark, Michael C. Jewett, Matthew P. DeLisa

## Abstract

Asparagine-linked (*N*-linked) protein glycosylation is one of the most abundant types of post-translational modification, occurring in all domains of life. The central enzyme in *N-*linked glycosylation is the oligosaccharyltransferase (OST), which catalyzes the covalent attachment of preassembled glycans to specific asparagine residues in target proteins. Whereas in higher eukaryotes the OST is comprised of eight different membrane proteins of which the catalytic subunit is STT3, in kinetoplastids and prokaryotes the OST is a monomeric enzyme bearing homology to STT3. Given their relative simplicity, these single-subunit OSTs (ssOSTs) have emerged as important targets for mechanistic dissection of poorly understood aspects of *N-*glycosylation and at the same time hold great potential for the biosynthesis of custom glycoproteins. To take advantage of this utility, this chapter describes a multipronged approach for studying and engineering ssOSTs that integrates *in vivo* screening technology with *in vitro* characterization methods, thereby creating a versatile and readily-adaptable pipeline for virtually any ssOST of interest.

Abbreviations

NLG: *N*-linked protein glycosylation
ssOST: single-subunit oligosaccharyltransferases
glycoSNAP: glycosylation of secreted *N*-linked acceptor proteins
CFPS: cell-free protein synthesis
IVG: *in vitro* glycosylation
LLOs: lipid-linked oligosaccharides
POPC: 1-palmitoyl-2-oleoyl-sn-glycero-3-phosphocholine

## 1. Introduction

Protein glycosylation is the attachment of glycans (mono-, oligo-, or polysaccharide) to specific amino acid residues in proteins, most commonly asparagine (*N*-linked) or serine and threonine (*O-*linked) residues. Roughly three-quarters of eukaryotic proteins and more than half of prokaryotic proteins are glycosylated (Dell, Galadari, Sastre, & Hitchen, 2010). Glycosylation adds an additional information layer to recipient proteins, modulating their folding and stability, receptor binding, enzymatic activity, and/or localization (Varki, 1993). Many glycoproteins reside on the cell surface where they influence myriad biological processes such as development (Haltiwanger & Lowe, 2004), innate and adaptive immunity (Daniels, Hogquist, & Jameson, 2002; Rudd, Elliott, Cresswell, Wilson, & Dwek, 2001), and host-microbe interactions in the gut (Tytgat & de Vos, 2016). Glycosylation also feature prominently in disease. For example, tumor cells commonly express glycans at atypical levels or with altered structural attributes (Lau & Dennis, 2008; Pinho & Reis, 2015) while many pathogens make use of glycans during invasion of host tissue (Benz & Schmidt, 2002; Valguarnera, Kinsella, & Feldman, 2016). Glycosylation is also vitally important to the development of many protein biologics, and has been harnessed for enhancing therapeutic properties such as half-life extension (Elliott et al., 2003; Flintegaard et al., 2010; Ilyushin et al., 2013; Lindhout et al., 2011), antibody-mediated cytotoxicity (Li et al., 2017; Lin et al., 2015), and immunogenicity (Lipinski et al., 2013; Sadoulet et al., 2007; Wacker et al., 2014). Yet despite the importance of glycosylation, forward progress in the field has lagged due in large part to a lack of tools for rapid and systematic characterization of the enzymes involved in the glycosylation process, in particular the oligosaccharyltransferase (OST). The net result is that glycans and their corresponding glycoconjugates remain one of the most important but least understood class of molecules in all of biology and medicine.

Eukaryotic and prokaryotic *N*-linked protein glycosylation (NLG) systems share many mechanistic features (Weerapana & Imperiali, 2006). Both involve enzymatic synthesis of a lipid-linked oligosaccharide (LLO) donor and transfer of the preassembled glycan from the lipid to the sequon of a target protein in a reaction that is catalyzed by an OST. In higher eukaryotes, the OST is comprised of eight different membrane proteins of which the catalytic subunit is STT3 (Yan & Lennarz, 2002), whereas in kinetoplastids (*i.e.*, *Trypanosoma brucei* and *Leishmania major*) and prokaryotes the OST is a monomeric enzyme bearing homology to STT3 (Lizak, Gerber, Numao, Aebi, & Locher, 2011; Matsumoto et al., 2013; Nasab, Schulz, Gamarro, Parodi, & Aebi, 2008). Amongst this latter group, the OST from the bacterium *Campylobacter jejuni*, named PglB (hereafter *Cj*PglB), has been most extensively studied and thus serves as the archetype for single-subunit OSTs (ssOSTs). Naturally, *Cj*PglB is part of a *bona fide* NLG pathway in *C. jejuni* where it catalyzes the *en bloc* transfer of preassembled heptasaccharide glycans from undecaprenol pyrophosphate (Und-PP) to protein substrates that bear the acceptor sequon D/E-X_-1_-N-X_+1_-S/T (where X_-1_ and X_+1_ ≠ P) (for a review, see ref. (Nothaft & Szymanski, 2010)). Shortly after its discovery, the entire *C. jejuni* NLG pathway including *Cj*PglB was functionally reconstituted in *Escherichia coli* (Wacker et al., 2002). Using this recombinant platform, it was demonstrated that PglB can transfer a wide array of structurally diverse oligosaccharides (Feldman et al.; Valderrama-Rincon et al.), highlighting its potential value in glycoengineering applications. However, while specificity towards the glycan donor is relaxed, *Cj*PglB recognizes a more stringent protein acceptor site compared to the N-X-S/T (X ≠ P) sequon recognized by eukaryotic OSTs (Kowarik, Young, et al., 2006). Specifically, *Cj*PglB requires an acidic residue in the -2 position of the sequon, thereby restricting bacterial NLG to a narrow set of polypeptides. To better understand this so-called “minus two rule” and whether it was conserved across Gram-negative bacterial species, we systematically characterized the acceptor site specificities of a diverse collection of more than 20 PglB homologs using an ectopic trans-complementation strategy in the same recombinant *E. coli* platform described above. Our metagenomic screening revealed that the majority of bacterial ssOSTs preferred a negatively charged residue in the -2 position, akin to *Cj*PglB; however, five ssOSTs were identified that recognized a broader range of acceptor sites (Ollis et al., 2015).

Directed evolution is emerging as an alternative strategy for shedding additional light on the sequence determinants governing the specificity of bacterial ssOSTs and for identifying unique ssOSTs with desirable substrate (i.e., glycan, protein acceptor) specificity with the potential to overcome some of the limitations of this system. The success of such directed evolution efforts hinges critically on the availability of high-throughput reporter assays that generate a genotype-glycophenotype linkage. To date, a handful of genetic screens for NLG have been described for this purpose, all of which combine glycoengineered *E. coli* carrying the complete protein glycosylation (*pgl*) locus of *C. jejuni* (Wacker et al., 2002) with a functional readout of glycosylation activity. Notable examples include: ELISA-based detection of periplasmic glycoproteins (Ihssen et al., 2012; Pandhal et al., 2013), glycophage display (Celik, Fisher, Guarino, Mansell, & DeLisa, 2010; Durr, Nothaft, Lizak, Glockshuber, & Aebi, 2010), cell surface display of glycoconjugates (Fisher et al., 2011; Mally et al., 2013; Valderrama-Rincon et al., 2012), and <u>glyco</u>sylation of <u>s</u>ecreted *<u>N</u>*-linked <u>a</u>cceptor <u>p</u>roteins (glycoSNAP) (Ollis, Zhang, Fisher, & DeLisa, 2014). Using this latter system, we successfully isolated *Cj*PglB variants recognizing the minimal N-X-S/T sequon used by eukaryotic OSTs (Ollis et al., 2014). One of the more interesting variants was capable of modifying a native sequon in the eukaryotic protein RNase A, an acceptor protein that had previously been inaccessible to the wild-type *Cj*PglB enzyme.

Here, we describe an integrated platform for studying and engineering ssOSTs that combines *in vivo* screening using glycoSNAP with state-of-the-art *in vitro* characterization methods, thereby creating a versatile and readily-adaptable pipeline for virtually any ssOST of interest (**Fig. 1**). The first step in the pipeline involves *in vivo* screening using glycoSNAP, whereby modified colony blotting on nitrocellulose membranes is used to create a genotype-glycophenotype linkage (**Fig. 2**). While our efforts to date have focused on identifying variants of bacterial ssOSTs that are able to modify non-canonical acceptor sequences, the assay could easily be extended to screen libraries of bacterial or non-bacterial ssOSTs (Matsumoto et al., 2013; Nasab et al., 2008) for variants that overcome other system limitations such as low efficiency with certain non-native glycan substrates (Ihssen et al., 2015). After screening, the identified ssOST candidates are subjected to further activity interrogation using an *in vitro* glycosylation (IVG) assay. The advantage of IVG is that it provides a platform where the glycosylation components can be easily decoupled and carefully investigated, in contrast to the more difficult to control NLG pathways in living cells. To facilitate rapid activity screening, candidate ssOSTs are prepared using a novel cell-free protein synthesis (CFPS)-nanodisc system that we recently developed (Schoborg et al., 2017). This latter system is capable of producing multiple active ssOSTs within a day, enabling facile and high-throughput activity screening of ssOSTs. Alternatively, ssOST candidates can be prepared using an economical alternative involving extraction of crude membrane extracts from *E. coli* cells expressing the ssOST of interest (Jervis et al., 2010). Finally, we provide optimized protocols for purification of active ssOSTs in high yield. Purified ssOSTs are valuable reagents for studying enzyme activity and mechanism under the most well-defined conditions. In addition, the purified enzyme is a prerequisite for structural interrogation using methods such as in-solution protein NMR (Huang, Mohanty, & Banerjee, 2010) and X-ray crystallography (Lizak et al., 2011; Matsumoto et al., 2013). Taken together, our comprehensive protocols provide a robust and modular pipeline for developing a suite of flexible, single-subunit *N-*glycosylation biocatalysts and growing the glycoengineering armament.

## 2. Materials

### 2.1 Media

1. Luria-Bertani (LB) broth: 1.0% (w/v) tryptone, 0.5% (w/v) yeast extract, 0.5% (w/v) NaCl, autoclave sterile.
2. LB agar: add 1.5% (w/v) agar to LB prepared as above. Aliquot 30 mL of LB agar per 150 mm petri dish into a plastic conical tube. Add appropriate antibiotics and sterile 0.2% (w/v) D-glucose. Mix and pour into petri dishes. For induction plates, omit glucose and add sterile 0.2% (w/v) L-arabinose and 0.1 mM isopropyl-β-D-thiogalactopyranoside (IPTG).
3. Terrific broth (TB): 1.2% (w/v) tryptone, 2.4% (w/v) yeast extract, 0.4% (v/v) glycerol, 10% (v/v) phosphate buffer (0.17 M KH_2_PO_4_, 0.72 M K_2_HPO_4_), autoclave sterile. Autoclave phosphate buffer separately from other components and add to the broth prior use.
4. 2xYTPG broth: 1.6% (w/v) tryptone, 1.0% (w/v) yeast extract, 0.5% (w/v) NaCl, 1.8% (w/v) glucose, 0.7% (w/v) K_2_HPO_4_, 0.3% (w/v) KH_2_PO_4_, adjust pH to 7.3 with 5 N KOH and autoclave sterile. Autoclave 40% (w/v) glucose stock separately and add to the broth prior use.

### 2.2 Media supplements

1. Transformation and storage solution (1×TSS): supplement LB broth with 10% (w/v) polyethylene glycol (PEG)-8000, 5% (v/v) dimethylsulfoxide (DMSO), and 20 mM MgSO_4_; adjust pH to 6.5 with HCl, and autoclave sterile.
2. Antibiotics: Ampicillin (Amp) is used at 100 μg/mL. To make a 1,000× stock, mix 1 g in 10 mL nanopure water. Chloramphenicol (Cam) is used at 20 μg/ mL. To make a 1,000× stock, dissolve 0.2 g in 10 mL ethanol. Trimethoprim (Tp) is used at 100 μg/mL. To make a 500× stock, dissolve 0.5 g in 10 mL DMSO. Kanamycin (Km) is used at 50 mg/mL. To make a 1,000× stock, dissolve 0.5 g in 10 mL nanopure water. All antibiotic stock is filter sterile.
3. Inducers/repressors: 20% (w/v) L-arabinose stock, 20% (w/v) D-glucose, and 0.1 M Isopropyl β-D-1-thiogalactopyranoside (IPTG) stock. All inducer/repressor stocks are made in nanopure water and filter sterilize.

### 2.3 Bacterial strains and plasmids

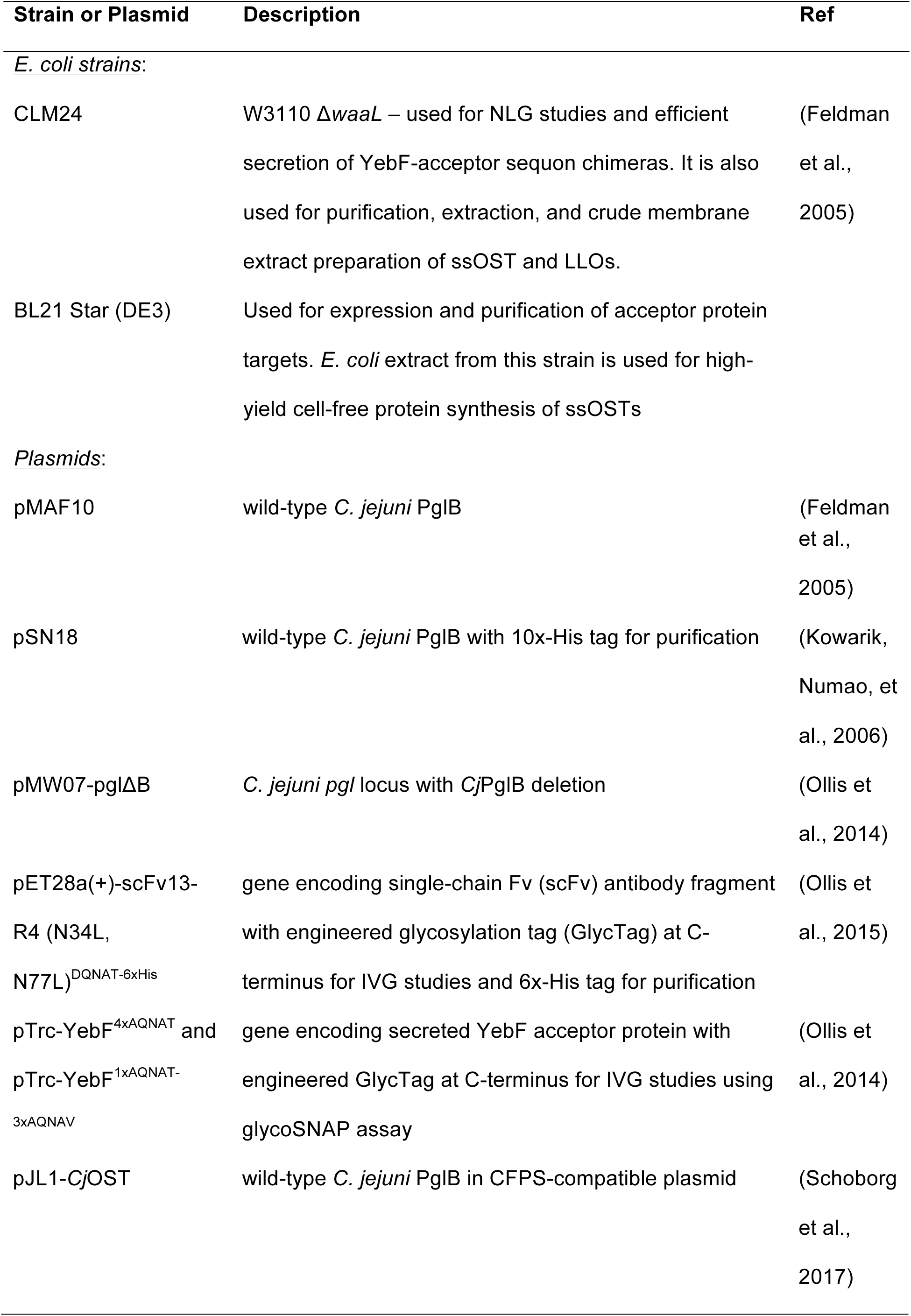

### 2.4 GlycoSNAP assay

1. Table-top centrifuge.
2. 0.45 μm, 142 mm Whatman cellulose nitrate filter membranes (VWR).
3. Nitrocellulose hybridization and transfer membranes (GE).
4. Sterile 1× phosphate buffered saline (PBS): 137 mM NaCl, 2.7 mM KCl, 10 mM Na_2_HPO_4_, and 1.8 mM KH_2_PO_4_ in nanopure water, autoclave sterile.
5. 30°C and 37°C stationary incubators.
6. Metal tweezers.
7. Flat-bottomed dish to fit membrane.
8. 20% (w/v) trichloroacetic acid (TCA) in nanopure water. <u>Caution!</u> Always wear gloves when handling TCA since it can cause severe burns Using nitrile gloves and handling TCA in fume hood are recommended.
9. Laemmli sample buffer: (for a 2× stock), mix 4 mL of 10% (w/v) sodium dodecyl sulfate (SDS), 2 mL of glycerol, 1.2 mL of 1 M Tris–HCl, pH 6.8, and 2.8 mL of nanopure water. Add 0.5 mg of bromophenol blue. Add β-mercaptoethanol to a final concentration of 5% (v/v).

### 2.5 Preparation of ssOSTs by CFPS

#### 2.5.1 S30 extract preparation

1. Avestin EmulsiFlex B15 (volumes < 15 mL) or C3 (>15 mL) high-pressure homogenizer.
2. High-speed centrifuge and rotor capable of spinning at 30,000xg.
3. 1× S30 extract buffer: 10 mM TrisOAc, 14 mM Mg(OAc)_2_, 60 mM KOAc, pH 8.2, filter sterile.
4. Sterile 50 mL falcon tube, 15 mL disposable conical tubes, and 1.5 mL microcentrifuge tube.
5. Aluminum foil.
6. Liquid nitrogen and dewar.

#### 2.5.2. Producing ssOST in CFPS supplemented with POPC-nanodiscs

1. S30 extract
2. Stock solutions: All stock solution is dissolved in nanopure water and filter sterile. The aliquots of the stocks are flash-frozen and stored at -80^o^C. Keep working stocks at -20^o^C for several months.
a. 15× salt solution (SS): 180 mM Mg(Glu)_2_, 150 mM NH_4_(Glu), and 1.950 M K(Glu)_2_.
b. 15× master mix (MM) stock: 18 mM adenosine triphosphate (ATP), 12.75 mM guanosine triphosphate (GTP), 12.75 mM uridine triphosphate (UTP), 12.75 mM cytidine triphosphate (CTP), 0.51 mg/mL folinic acid, and 2.559 mg/mL *E. coli* tRNA (Roche).
c. 15× reagent mix (RM) stock: 50 mM amino acids mix, 1 M phosphoenolpyruvate (PEP, Roche), 100 mM nicotinamide adenine dinucleotide (NAD), 50 mM coenzyme-A (CoA), 1 M oxalic acid, 250 mM putrescine, 250 mM spermidine, and 1 M HEPES.
3. Purified 1-palmitoyl-2-oleoyl-sn-glycero-3-phosphocholine (POPC) nanodiscs at 15 mg/mL stock concentration, prepared according to the standard protocol (Bayburt, Grinkova, & Sligar, 2002).
4. Nuclease-free water.
5. Sterile 1.5 mL microcentrifuge tube.

### 2.6 Protein purification and crude membrane extracts containing ssOST or LLOs

1. Buffer A Resuspend buffer: 50 mM HEPES, 250 mM NaCl, pH 7.5, filter sterile with 0.2 μm bottle-top filter.
2. Pierce™ Protease Inhibitor Tablets, EDTA-free.
3. RNase-Free DNase I from EpiCentre.
4. Avestin EmulsiFlex C5 homogenizer.
5. Buffer B: buffer A supplied with 10% (v/v) Glycerol and 1% (w/v) n-Dodecyl β-D-maltoside (DDM), at pH 7.5 and filter sterile.
6. Centrifuge and ultracentrifuge with rotor capable of spinning at 100,000xg.
7. Potter-Elvehjem tissue homogenizer.
8. Ni-NTA affinity resin.
9. Gravity column for affinity purification.
10. Nickel affinity purification buffers:

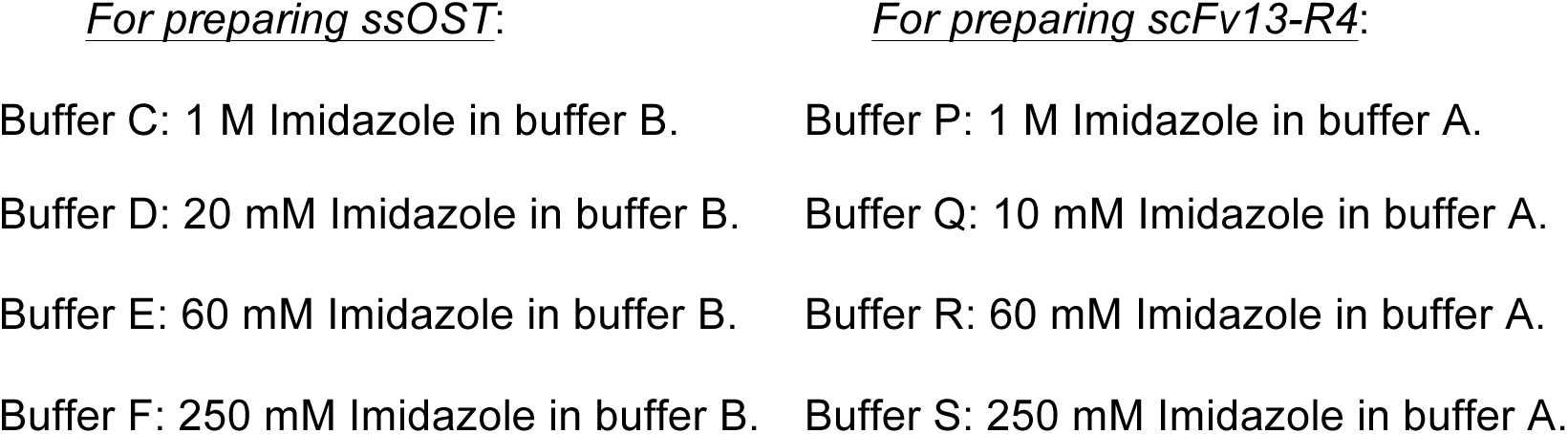
11. ÄKTA fast protein purification (FPLC) system with SuperDex-200 size exclusion chromatography column.
12. Size exclusion buffers using with ÄKTA system:

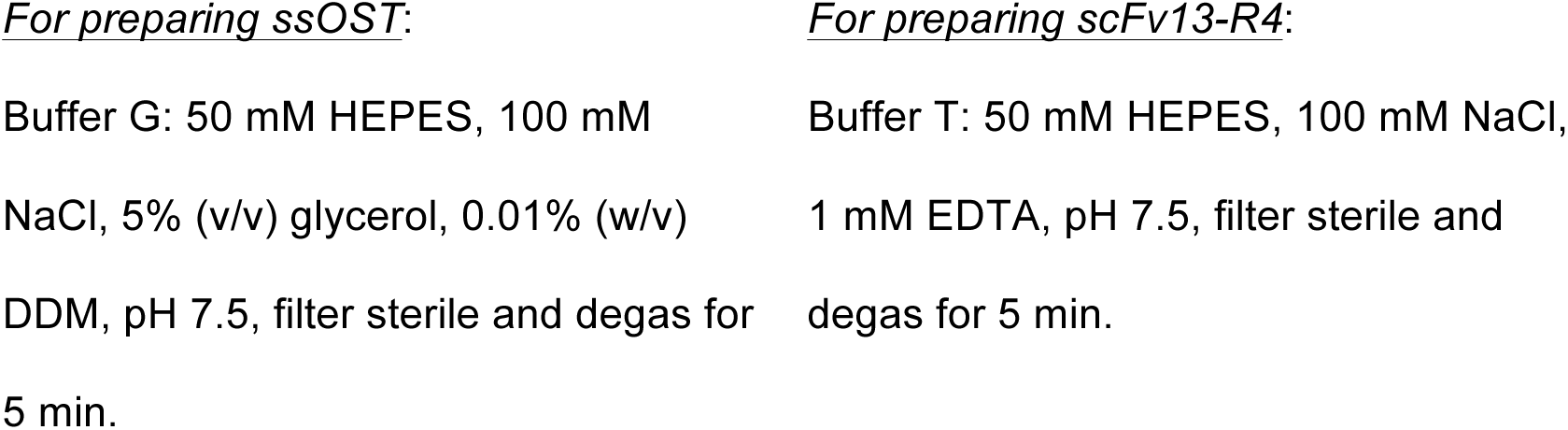
13. 3K MWCO protein concentrator column.
14. BioRad Bradford protein concentration assay.
15. BioRad RC DC™ (reducing agent and detergent compatible) protein concentration assay.

### 2.7 Extraction of LLOs

1. Lyophilizer.
2. 30 mL PTFE-conical tube.
3. Extracting solution: 10:20:3 (v/v/v) chloroform: methanol:water. Measure and mix all solvent in glassware since chloroform will dissolve plastic. Store the solution in a capped bottle at all time to prevent concentration change due to evaporation of chloroform and methanol. <u>Caution!</u> Always wear gloves and use fume hood when handling the extracting solution since chloroform is toxic.
4. Clean metal spatula.
5. 15 mL clean glass vial.
6. Vacuum concentrator machine that withstand organic solvent.

### 2.8 IVG reaction

1. Sterile 1.5 mL microcentrifuge tube.
2. 10× IVG buffer stock: 100 mM HEPES, 100 mM MnCl_2_, 1% (w/v) DDM, pH 7.5 in nanopure water and filter sterile.
3. Sterile nanopure water.
4. 30^o^C stationary water bath.

### 2.9 Lectin blot and Western blot analysis of glycosylation products

1. Standard apparatus for SDS-PAGE and immunoblotting analysis.
2. Immobilon-P PVDF 0.45μm membrane.
3. Tris-buffered saline (TBS): dissolve 80.0 g of NaCl, 20.0 g of KCl, and 30.0 g of Tris base in 800 mL of nanopure water, bring volume to 1 L and autoclave.
4. Tris-buffered saline, 0.05 % Tween-20 (TBST): Add 100 mL of 10× TBS to 900 mL of nanopure water. Add 500 μL Tween-20.
5. Albumin from bovine serum (BSA): 5% (w/v) in TBST for blocking solution, 3% (w/v) in TBST for lectin blotting. When using lectin, BSA/TBST is a preferred blocking solution than milk/TBST to prevent interaction between lectin and milk oligosaccharides.
6. Non-fat dry milk Nestle^®^: 5% (w/v) in TBST for blocking solution for all other antibodies.
7. Anti-6xHis Tag antibody peroxidase conjugate (His-HRP, from Abcam): 0.5 μg/mL in 5% milk/TBST.
8. Soybean agglutinin peroxidase conjugate (SBA-HRP): 0.5 μg/mL in 1% BSA/TBST (or other lectin or antibody specific for the glycan of choice).
9. Rabbit serum containing anti-*C jejuni* heptasaccharide glycan antibody (HR6P-Rabbit) [Note 1]: 0.5 μg/mL in 5% milk/TBST.
10. Goat Anti-Rabbit IgG H&L peroxidase conjugate (Rabbit-HRP, from Abcam): 0.05 μg/mL in 5% milk/TBST.
11. BioRad Clarity™ Western ECL Substrate.
12. % Coomassie Brilliant Blue membrane stain solution: dissolve 0.1 g of Coomassie Blue R250 in 50 mL of methanol (MeOH), 7 mL of acetic acid, and 43 mL of nanopure water. Stain can be saved and reused multiple times.
13. Destain solution: 50 % MeOH in nanopure water. Discard destain following hazardous waste protocols.
14. BioRad ChemiDoc™ XRS+ System.

## 3. Methods

### 3.1 GlycoSNAP assay

Days 0-1 Transformation of glycocompetent *E. coli* for library screening

1. Inoculate 5 mL LB supplemented with 0.2 % D-glucose and antibiotics as needed with a single colony of the strain to be transformed. Grow overnight at 37 °C.
2. Subculture 1:100 from the overnight culture into a fresh 5-mL volume of the same medium. Grow until culture density (OD_600_) reaches 0.4–0.5.
3. Harvest 5 × 1 mL into Eppendorf tubes [Note 2]. Chill on ice for 5 min. Pellet cells at 4°C in a tabletop centrifuge. Discard supernatant. Resuspend cell pellets in 100 μL of ice-cold 1× TSS.
4. Add 50-200 ng of library plasmid miniprep to the prepared cells. Incubate on ice for 30 min. Heat shock 90s at 42 °C. Immediately add 500 μL of LB to rescue cells. Incubate for 1 h at 37 °C with aeration. Pour LB agar plates as needed in preparation for the next step.
5. Plate at least 100 μL of cells and spread evenly using a spreader or sterile beads. A plating with optimal cell density for screening should yield about 2,500 colonies on a 150-mm plate. Incubate plates at 37°C overnight.

Days 2-3 GlycoSNAP Assay

6. Trim one cellulose nitrate filter circle and one piece of nitrocellulose membrane to fit a 150-mm plate (one set for each transformation plate to be screened). Cut two notches on both filter and membrane to assist in later alignment.
7. Pre-wet the nitrocellulose membrane in 1× PBS, keeping the matte side up, and place onto a fresh induction plate. Cover with lid to prevent drying in between steps.
8. Replicate colonies from transformation plate by gently placing the cut filter membrane directly onto the plate to avoid air bubbles. The side in contact with the colonies should be the side that was not in contact with the nitrocellulose when stacked to cut.
9. Using sterilized metal tweezers, carefully peel up the colony-containing membrane and place colony-side-up onto the nitrocellulose membrane on the induction plate. Ideally, match notches on the filter and nitrocellulose membrane.
10. Incubate plates right-side-up at 30°C overnight (16-18 h).
11. The next day, use tweezers to remove the colony-containing membrane and transfer it onto a fresh LB agar plate. Save at 4°C. Transfer the nitrocellulose membrane into a dish of 1× TBS. Shake at room temperature about 10 min to rinse [Note 3].
12. Block membrane for 1 h in 5% BSA/TBST blocking solution.
13. Incubate for 1 h with SBA-HRP solution. 30 mL of solution is sufficient to cover the ~140-mm membrane circle.
14. Wash 4× with TBST, incubating each wash at least 10 min with shaking.
15. Develop blot.
16. If desired, the blot can be stripped with standard Western blot stripping buffer and reprobed using antibodies specific for the secreted target (anti-His for the C-terminal 6xHis tag fused to the YebF construct) and/or the membrane can be stained with a general protein stain such as Coomassie blue.

Days 3-5 Confirmation of positive hits

17. Pick individual colonies identified as positive hits and restreak on LB agar plates containing the appropriate antibiotics. Incubate at 37°C overnight.
18. Inoculate a single colony into 5 mL of LB supplemented with 0.2% D-glucose and appropriate antibiotics for each hit to be tested and grow overnight at 37°C. A control such as a strain expressing the wild-type *C. jejuni pgl* locus and YebF DQNAT should be included for comparison.
19. The next day, subculture the overnight cultures 1:100 in 5 mL of LB supplemented with the appropriate antibiotics and grow at 37°C to an OD_600_ ~ 0.6. Induce with 100 μM IPTG and 0.2 % L-arabinose. Incubate at 30°C overnight.
20. The next day, harvest 1 mL of each culture and pellet cells 5 min at 4°C. Determine protein concentration in the supernatants using a Bradford assay. Harvest volumes with equal protein concentrations and precipitate protein by addition of an equal volume of ice-cold 20% TCA. Vortex and incubate on ice for at least 15 min. Pellet precipitated protein by centrifuging at 10,000×g, 5 min, at 4°C. Discard supernatant. Centrifuge briefly a second time and remove any residual acid. Resuspend pellets in 25 μL of 1 M Tris–HCl, pH 7.5 then add 25 μL of 2× Laemmli sample buffer. Boil 5- 10 min.
21. Detect glycosylation state by standard SDS-PAGE and immunoblotting (see section 3.5).
22. Plasmids from true positive hits can be isolated and sequenced to identify mutations conferring activity.

### 3.2 Preparation of ssOST by CFPS

#### 3.2.1 S30 extract preparation

Days 0-2

1. Grow *E. coli* strain BL21 Star (DE3) in a shake flask or fermenter to OD_600_ ~3 in 2×YTPG media. Add 1 mM IPTG at OD_600_ 0.6-0.8 to induce expression of T7 RNA polymerase.
2. Harvest cells by centrifugation at 5000xg for 15 min at 4°C. Wash cells 3x in 25 mL of S30 buffer (vortex to resuspend) and pellet by centrifugation at 5000xg for 10 min at 4°C. After the last resuspension, pellet cells at 8000×g for 10 min at 4°C and flash-freeze on liquid nitrogen. Pellet can be stored at -80°C or used directly.

Day 3 preparing extract

*Keep the extract on ice at all times unless noted otherwise. Work seamlessly. All equipment in contact with lysate should be pre-equilibrated to 4*°*C*.

3. Remove cell pellet from -80°C and add 1 mL of S30 buffer per 1 g of wet cell mass. Dislodge pellet from the wall of the bottle. Vortex to resuspend to homogeneity.
4. Disrupt cells using Avestin EmulsiFlex-B15 (lysis volumes <15 mL) or C3 (>15 mL) high-pressure homogenizer at 20,000-25,000 psi. Pass the cells only once. Cell lysis is the key step in extract preparation and could be alternatively performed using sonication (Kwon & Jewett, 2015).
5. Centrifuge lysate at 30,000×g for 30 min at 4°C to remove cell debris.
6. Immediately pipette the supernatant into new centrifuge tubes and centrifuge again at the same setting.
7. Immediately pipette supernatant into 1.5-mL microcentrifuge tubes.
8. Pre-incubation: wrap the microcentrifuge tubes in aluminum foil and incubate at 37°C in a shaker (~120-250 rpm) for 60 min.
9. Clarification: centrifuge at 15,000×g for 15 min at 4°C. Immediately pipette the supernatant into 15 mL disposable conical tubes and place on ice.
10. Immediately make 50 μL aliquot and 1-2 mL volume stocks of the cell extract. Flash freeze in liquid nitrogen and store at -80°C.
11. Perform a Bradford assay to measure total protein concentration (usually ~40 g/L). S30 extract performance is maintained for approximately 3 freeze-thaw cycles.

#### 3.2.2 Producing ssOST in CFPS supplied with POPC-nanodiscs

1. Calculate the appropriate volumes of each reagent according to total number of reactions. Ensure that all components are at the concentrations listed.
2. Thaw all the reagents on ice. Set up microcentrifuge tubes on ice, one for each cell-free reaction and one for reaction premix.
3. CFPS reaction is performed with a modified, reducing PANOx-SP system (Jewett & Swartz, 2004):

| CFPS components | Volume per 1 reaction (µL) |
| --- | --- |
| 15x SS | 1 |
| 15x MM | 1 |
| 15x RM | 1 |
| pJL1 plasmid encoding ssOST | 200 ng |
| S30 extract | 4 |
| 15 mg/mL POPC nanodiscs | 1 |
| Nuclease-free water | Bring final volume to 15 µL |

a. To make a premix, combine the appropriate amounts of 15×SS, 15×MM, 15×RM, plasmid and nuclease-free water in reaction premix tube. Vortex and quickly spin down the tube [Note 4].
b. Then add the appropriate amounts of POPC nanodiscs and S30 extract.
c. Gently pipette the mixture up and down to thoroughly mix all components, but make sure to minimize bubble formation.
4. Aliquot 15 μL premix into individual microcentrifuge tube.
5. Briefly centrifuge CFPS-reaction tubes to ensure all liquid is held at the bottom of the tube.
6. Incubate reaction at 30°C for 6 h in a stationary water bath. The CFPS reaction containing ssOST in nanodisc can be loaded directly into IVG reaction for rapid glycosylation screening.

### 3.3 Preparation of purified and crude membrane extract glycosylation components

#### 3.3.1 Preparation of crude membrane extract or purified ssOST enzyme

Days 0-1

1. Grow *E. coli* strain CLM24 carrying pSN18 plasmid in 50 mL LB supplied with ampicillin and 0.2% D-glucose overnight at 37°C.
2. Subculture 1:20 from the overnight culture into a fresh 1.0 L TB supplied with ampicillin. Grow with shaking at 220 rpm until OD_600_ reaches 0.4–0.5.
3. Adjust incubation temperature to 16^o^C and leave the culture with shaking for an hour.
4. Induce protein expression with 0.02% L-arabinose. Incubate at 16°C overnight.

Day 2 prepare membrane extract containing active *Cj*OST

5. The next day, harvest cell by centrifugation at 8,000×g for 10 min at 4°C. Wash cell pellet by resuspending with 200 mL buffer A and centrifuge again at the same setting. Discard supernatant. Collect pellet and determine wet cell mass. Pellet can be saved in -80°C fridge for a month or used directly.
6. Resuspend cell pellet using 10 mL buffer A per 1 g wet cell mass. Add EDTA-free protease inhibitor to prevent protein degradation. Add DNAse to reduce sample viscosity. Use standard manufacturer’s protocol.
7. Pre-equilibrate Avestin homogenizer with ice-cold buffer A. Disrupt cells using Avestin C5 EmulsiFlex homogenizer at 17,000 psi for 3 passes.
8. Centrifuge lysate at 30,000×g for 30 min at 4°C to remove cell debris.
9. Collect and ultracentrifuge supernatant at 100,000×g for 2 h at 4°C to isolate membrane fraction.
10. Collect pellet containing membrane fraction and *Cj*OST. Resuspend pellet in 20 mL buffer B using Potter-Elvehjem tissue homogenizer. Make sure to fully resuspend the pellet [Note 5]. Transfer homogenized sample into sterile 50 mL conical tube. Add protease inhibitor cocktail into sample and incubate with shaking (120 rpm) at room temperature for an hour. The DDM detergent in buffer B will extract and solubilize *Cj*OST from bacterial membrane.
11. Ultracentrifuge sample at 100,000×g for an hour at 4°C. The supernatant now contains detergent-solubilized *Cj*OST enzyme. <u>Alternatively:</u> To prepare crude membrane extract containing active *Cj*OST, centrifuge sample at 20,000×g for an hour at 4°C. Collect and immediately add protease inhibitor into supernatant after centrifugation. The crude membrane extract is active at 4°C for one week. We have demonstrated the use of this method to prepare several active ssOSTs that can modifty targeted protein acceptor. In addition, crude membrane extract containing active LLOs can be prepared in a similar method [Note 6].
12. Add 0.4 mL buffer C into supernatant to adjust imidazole concentration to 20 mM.
13. Equilibrate 0.5 mL Ni-NTA resin by washing with ice-cold buffer D at 5 times bed volume. Add pre-equilibrated Ni-NTA resin into supernatant, incubate with rolling overnight at 4°C.

Day 3 purification by affinity and size exclusion chromatography (SEC)

*Keep sample on ice at all times unless noted otherwise. All equipment in contact with sample should be pre-equilibrated to 4*°*C*.

14. Load sample into clean gravity column at the flowrate of 0.5 mL/min.
15. Wash resin with 5 bed volumes of buffer D, followed by 5 bed volumes of buffer E. Then elude protein with 7 bed volumes of buffer F. Keep all the fractions for analysis by Coomasie blue.
16. Pre-equilibrate SuperDex-200 SEC column connecting ÄKTA-FPLC system with ice-cold buffer G. Load eluent fraction into sample loop. Inject sample through SEC column. Collect and combine fractions with size corresponding to *Cj*OST (84 kDa) together.
17. Concentrate protein to 1-2 mg/mL final concentration using 3K MWCO protein concentrator column. Add glycerol to the sample at 20% (v/v) concentration. Aliquot and store *Cj*OST at -80°C for 4-5 months.
18. Determine protein concentration and sample purity with RC/DC assay and Coomassie blue protein stain, respectively.

#### 3.3.2 Purification of acceptor protein scFv13-R4(N34L, N77L)^DQNAT-6xHis^

1-8 These steps are essentially the same as protocol described in section 3.2.1 above, with a few exceptions. *E. coli* strain BL21 Star (DE3) carrying pET28a(+)-scFv13- R4(N34L, N77L)^DQNAT-6xHis^ plasmid is used. The inducer for pET-based vector is IPTG at 0.1 mM final concentration. Kanamycin antibiotic is used at 100 ug/mL.
9. Adjust the imidazole concentration in the supernatant to 10 mM Imidazole with buffer P.
10. Equilibrate 0.25 mL Ni-NTA resin by washing with ice-cold buffer Q at 5 times bed volume. Add pre-equilibrated Ni-NTA resin into supernatant, incubate with rolling at room temperature for an hour.

*Keep sample on ice at all times unless noted otherwise. All equipment in contact with sample should be pre-equilibrated to 4*°*C*.

11. Load sample into clean gravity column at the flowrate of 0.5 mL/min.
12. Wash resin with 5 bed volumes of buffer Q, followed by 5 bed volumes of buffer R. Then elude protein with 7 bed volumes of buffer S. Keep all the fractions for analysis by Coomasie blue.
13. Pre-equilibrate SuperDex-200-SEC column connecting ÄKTA-FPLC system with ice-cold buffer T. Load eluent fraction into sample loop. Inject sample through SEC column. Collect and combine fractions with size corresponding to scFv13-R4 (29 kDa) together.
14. Concentrate protein to 1-2 mg/mL final concentration using 3K MWCO protein concentrator column. Add glycerol to the sample at 10% (v/v) concentration. Aliquot and store scFv13-R4 at -80°C for 6 months.
15. Determine protein concentration and sample purity with Bradford assay and Coomassie blue protein stain, respectively.

#### 3.3.3 Extraction of LLOs bearing *C. jejuni* glycan (adapted from (Guarino & DeLisa, 2012))

Days 0-2

1-5 These steps are essentially the same as protocol described in section 3.2.1, with a few exceptions. *E. coli* strain CLM24 carrying pMW07-pglΔB plasmid is used. The inducer for this plasmid is L-arabinose at 0.2% final concentration. Chloramphenicol antibiotic is used at 20 ug/mL.
6. Use clean spatula to scrap cell pellet and transfer to clean 50 mL conical tubes. Freeze-dry cell pellets to complete dryness at -70°C with lyophilizer (usually takes ~2 days).

Day 4

7. Weigh and combine lyophilisate into a sterile 30 mL PTFE-conical tube. Use clean spatula to break dried pellet into small fractures.
8. Add 20 mL 10:20:3 (v/v/v) chloroform:methanol:water extracting solution into the tube and incubate with shaking for 30 min at room temperature.
9. Centrifuge the mixture at 4000×g for 15 min at 4°C.
10. Transfer organic fraction (bottom layer) to a clean 15 mL glass vial. Remove chloroform and methanol with vacuum concentrator at room temperature (usually take ~4-5 h).
11. Place the vial into freeze-dry unit to remove residue water at -70°C overnight.

Day 5

12. Lyophilisate now contains active lipid-linked oligosaccharide (*Cj*LLOs). Weigh lyophilisate mass. Dried LLOs can be stored at -80°C for 6 months.
13. Resuspend lyophilisate at 1.0 mL 1× IVG buffer per 1.0 mg lyophilisate dried weight. The resuspension should look yellowish. Transfer the mixture to a sterile microcentrifuge tube, spin down briefly, aliquot and store soluble fraction containing active *Cj*LLOs in -20°C for up to 2 months.

### 3.4 IVG setup

Day 1

1. In a sterile 1.5-mL microcentrifuge tube, add following reagents:

a. 3 μg purified antibody fragment scFv13-R4. Alternatively, N-terminal TAMRA-labelled peptide at 8.5 μM can be used as an acceptor substrate [Note 7].
b. 2 μg purified *Cj*OST or 25 μL crude membrane extract containing active *Cj*OST or 25 μL CFPS-nanodisc reaction containing active *Cj*OST.
c. 5 μg extracted *Cj*LLOs.
d. 5 μL 10× IVG buffer.
e. Bring final volume to 50 μL with sterile nanopure water.
2. In addition, it is necessary to set up control reactions to prevent fault-positive result. A typical reaction set is as follow:

3. Incubate the reaction tube in stationary water bath at 30°C for 16 h.

| IVG components | Sample | Control |  |  |
| --- | --- | --- | --- | --- |
| Protein/peptide acceptor | + | + | + | - |
| CjOST | + | + | - | + |
| CjLLOs | + | - | + | + |

Day 2

4. Centrifuge reaction tube at 10,000×g for 15 min at 4°C.
5. Collect soluble fraction. Reaction is stopped by adding Laemmli sample buffer. Keep sample at -20°C for analysis by SDS-PAGE followed by immunoblotting.

### 3.5 Detection of glycoprotein from *in vivo* and *in vitro* glycosylation assay

1. Load sample containing 50 μg total protein from *in vivo* experiment (see section 3.1) or 0.5 μg acceptor protein from IVG (see section 3.4) into SDS-polyacrylamide gels. Run protein electrophoresis at 200 V for 45 min [Note 8]. <u>Alternatively:</u> To detect fluorophore labelled-glycopeptide, load 10 μL sample into tris-tricine polyacrylamide gel. The electrophoretic condition is at 30 V initial voltage for 1 h and then 190 V for 3 h. The peptide can be visualized in-gel using any fluorescence imager.
2. Transfer protein sample onto two PVDF membrane. After transfer, wash membranes briefly with 10 mL 1× TBS buffer.
3. Incubate membranes with milk/TBST blocking solution for an hour at room temperature. BSA/TBST blocking solution is used instead for lectin blotting.
4. Wash membrane 4× with TBST, incubating each wash at least 10 min with shaking.
5. Immunoblot one membrane with anti-His antibodies and one with glycan-specific lectin (SBA-HRP) or antibodies (hR6P) for an hour at room temperature. Then wash membrane 4× with TBST, incubating each wash at least 10 min with shaking. The anti-His and lectin SBA-HRP immunoblotting membranes are ready to develop.
6. Incubate hR6P immunoblotting membrane with anti-rabbit-HRP secondary antibody for an hour at room temperature. Then wash membrane 4× with TBST, incubating each wash at least 10 min with shaking. The anti-glycan immunoblotting membrane is ready to develop.
7. To develop immunoblotting membrane, apply 1.0 mL of Western ECL substrate per membrane (9 x 7 cm). Incubate 5 min with shaking at room temperature. Use ChemiDoc™ XRS+ System to scan chemiluminescent signal.

## 4. Conclusion

Bacterial ssOSTs are highly modular enzymes, but we have only scratched the surface on exploiting their full biocatalytic potential, including identification of mutant activities. The protocols described here provide a robust framework for (1) understanding naturally occurring ssOSTs found in the genomes of kinetoplastid and prokaryotic organisms and (2) identifying entirely novel ssOSTs with desired glycophenotypes such as specificity for target acceptor sequons and/or glycan structures. The development and application of CFPS-nanodiscs and IVG assays provides a complementary set of techniques for synthesizing ssOSTs and subsequently evaluating their activity, all within a couple of days and with minimal technical difficulty. Finally, the optimized protocol for high-yield preparation of purified ssOST enzymes will facilitate thorough biochemical and structural analysis. Overall, our pipeline is expected to extend the glycoengineering toolkit for the facile discovery of novel glycosylation biocatalysts with customized functions.

**Figure 1.**
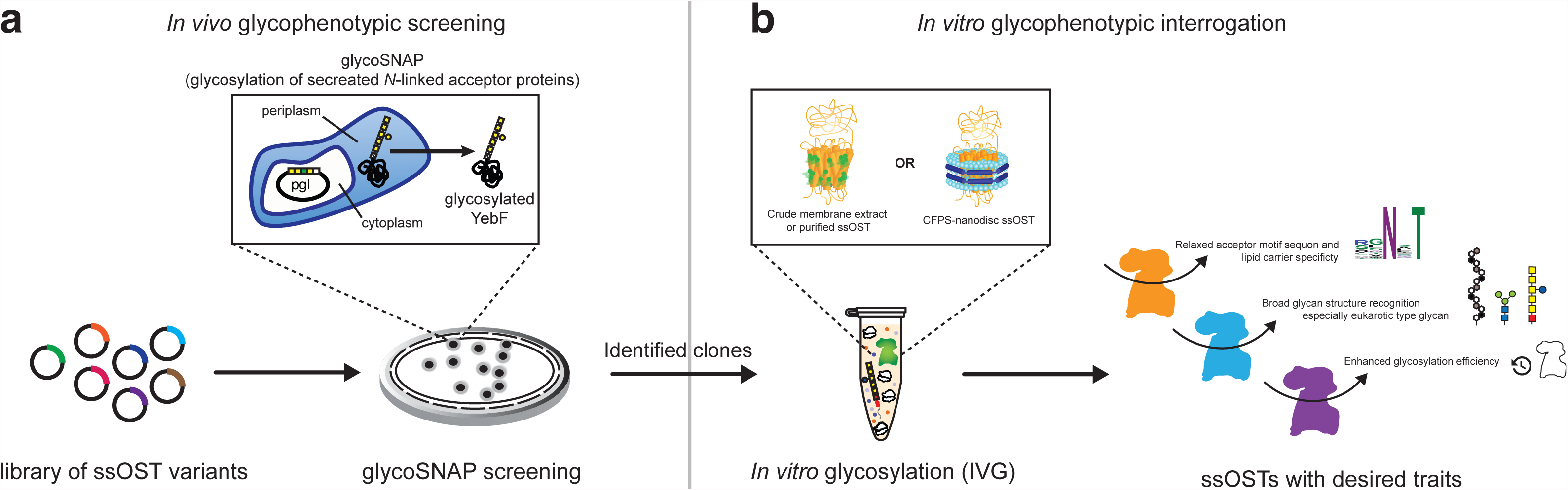
An integrated pipeline for studying and engineering ssOSTs. **(a)** Glycocompetent *E. coli* strain carrying YebF acceptor protein and glycan biosynthesis pathway (*e.g.*, *pgl* locus) is transformed with a combinatorial library of ssOST variants. The library is screened using the glycoSNAP assay, a high-throughput screening methodology using modified colony blotting to generate a genotype-glycophenotype linkage. **(b)** The isolated variants are then subjected to *in vitro* production and characterization methods with a goal of developing detailed structure-activity relationships (SARs) for each ssOST. Functional analysis under tightly controlled conditions is performed using IVG with purified protein acceptor, extracted LLOs as the glycan donor, and one of the following: crude membrane extract containing active ssOST enzyme, purified ssOST enzyme, or CFPS-derived ssOST supplemented with POPC-nanodiscs. Protein glycosylation is confirmed by SDS-PAGE and immunoblotting with glycan-specific antibody or lectin. The entire process takes only about two weeks and yields a set of ssOST variants with desired traits.

**Figure 2.**
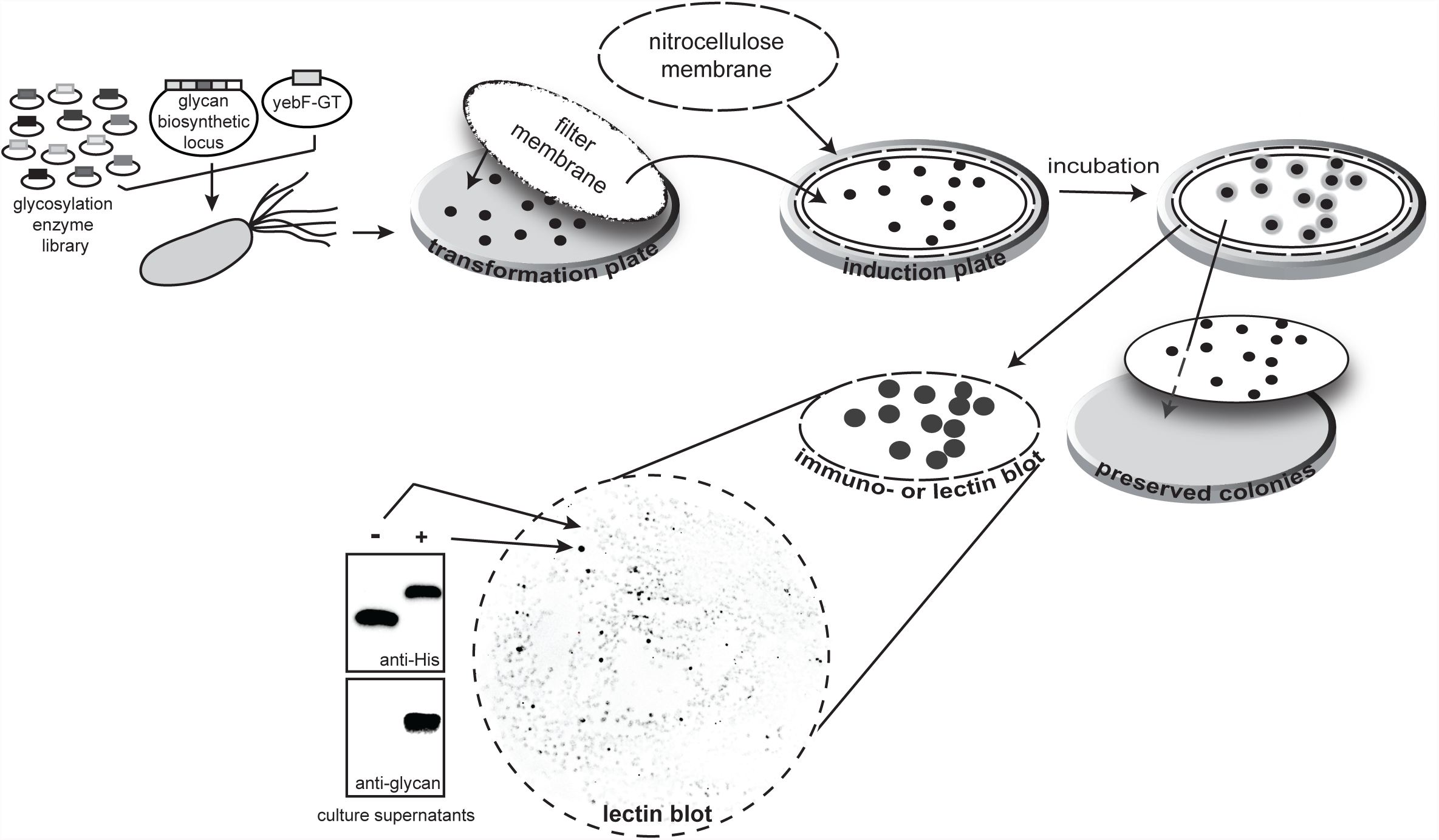
The glycoSNAP screening methodology. Glycocompetent *E. coli* strain carrying YebF acceptor protein and glycan biosynthesis locus is transformed with a combinatorial library of ssOST variants and plated on agar plate(s). Filter membrane is used to replicate colonies onto induction plate(s) containing inducers for the expression of YebF and NLG pathway enzymes. The glycosylated YebF is secreted out of the cells, after which it is immobilized on overlaid nitrocellulose membranes and detected by subjecting the nitrocellulose membrane to immunoblotting with glycan-specific antibody or lectin. Glycosylation competent (+) and incompetent (-) clones are verified by Western blotting of liquid culture supernatants using anti-His antibodies to detect the acceptor protein and anti-glycan antibodies (or lectins) to detect the oligosaccharide.

**Figure 3.**
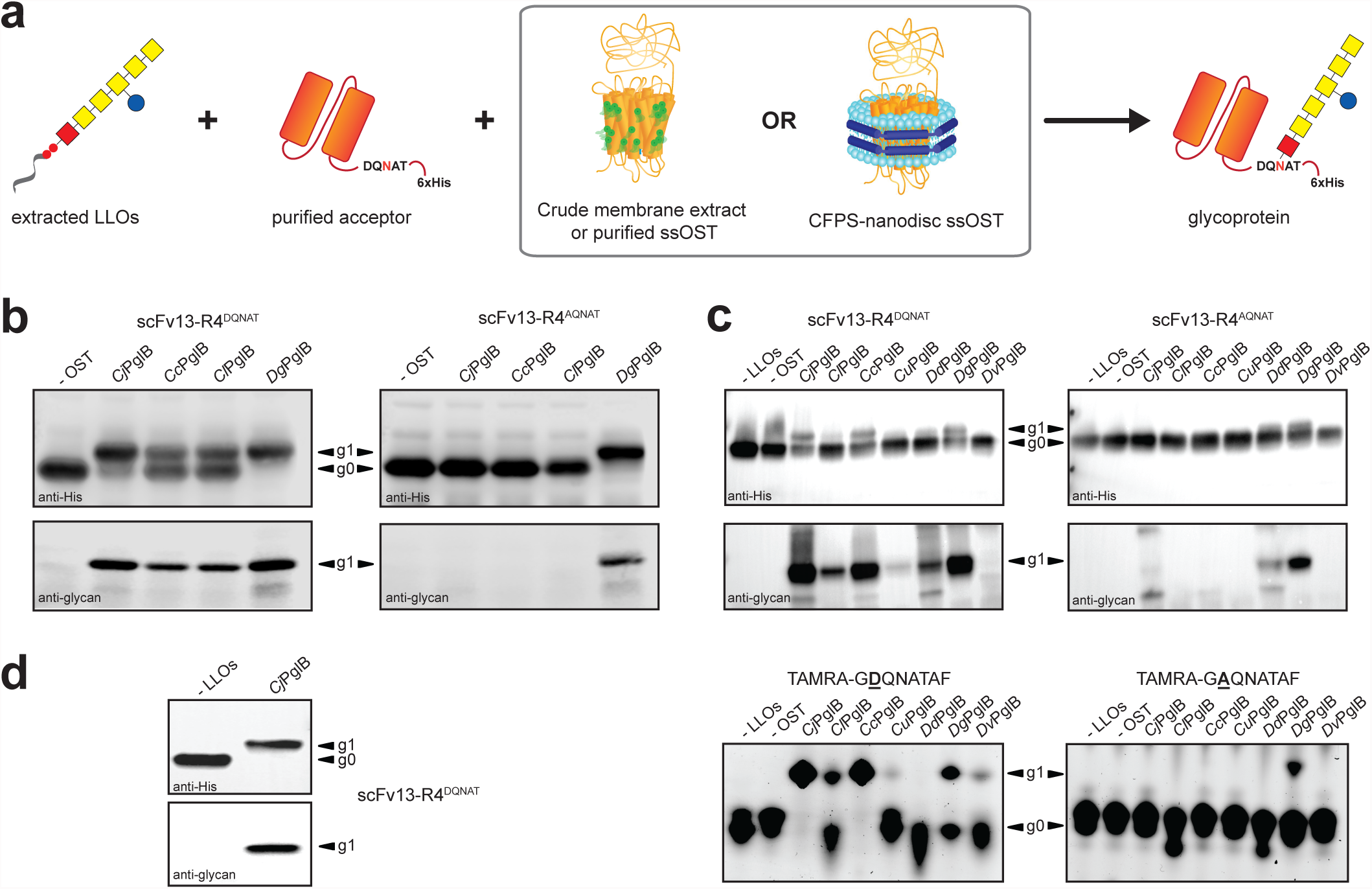
Cell-free production and characterization of ssOSTs. **(a)** Schematic of IVG for *in vitro* modification of purified acceptor peptides/proteins in the presence of extracted LLOs and ssOSTs that are provided by one of the following methods: crude membrane extraction or purification from cells expressing ssOST enzyme, or CFPS-based expression in the presence of POPC-nanodiscs. **(b)** Immunoblot analysis of IVG products generated by incubating the following: (1) LLOs, (2) POPC-nanodiscs containing bacterial PglB homologs from *C. jejuni* (*Cj*PglB), *C. coli* (*Cc*PglB), *C. lari* (*Cl*PglB) and *Desulfovibrio gigas* (*Dg*PglB), and (3) the acceptor protein scFv13-R4 containing a C-terminal glycosylation tag (GlycTag) encoding either a canonical (scFv13-R4^DQNAT^) or non-canonical (scFv13-R4^AQNAT^) sequon. **(c)** Similar analysis as in (b) using bacterial PglB homologs prepared in crude membrane extracts (top panels). Additional homologs included PglB from *C. upsaliensis* (*Cu*PglB), *D. desulfuricans* (*Dd*PglB), and *D. vulgaris* (*Dv*PglB). IVG of TAMRA-labeled peptides by extracted LLOs and crude membrane extracts containing bacterial PglB homologs (bottom panels). IVG-derived products were resolved by tricine/SDS-PAGE, and fluorescence signals were acquired with an image analyzer. **(d)** Similar analysis as in (b) using purified *Cj*PglB. In all blots, negative control reactions lacking LLOs (- LLOs) and/or ssOSTs (- OST) were included. Arrows denote aglycosylated (g0) and singly glycosylated (g1) forms of the scFv13-R4^DQNAT^ protein or TAMRA peptides. Blots were probed with antibody against the C-terminal 6xHis tag (anti-His) on the acceptor protein or anti-glycan serum reactive with *C. jejuni* heptasaccharide (anti-glycan).

## Notes

1. Rabbit serum containing *C jejuni* heptasaccharide glycan-specific antibody (hR6P) is made in-house and generously provided from Prof. Markus Aebi at ETH-Zürich, Switzerland.

2. Usually at least two transformations were done for each set to allow for plating of different cell densities or to ensure at least one plate with even spreading (the best one or both can be chosen to proceed with the assay).

3. This rinsing step was found to be important for cleaner blots so do not skip. For all blotting steps, it is important that the shaking evenly covers the membrane with the buffers. Insufficient shaking will result in uneven signal that will make it difficult to pick positive hits.

4. If different targeted ssOSTs will be produced in CFPS-nanodisc reaction, omit plasmid in premix and add each plasmid into individual reaction later.

5. Solubilization is a critical step in extracting active ssOST from the *E. coli* membrane. It is important to completely homogenize the sample and allow sufficient incubation time with DDM detergent to maximize extracting efficiency.

6. Similarly, crude membrane extract LLOs can be prepared the same way. Prepare 1 L TB culture with *E. coli* strain CLM24 carrying pMW07-pglΔB plasmid. After protein expression and cell harvesting, disrupt cell with Emulsiflex C5 homogenizer. Ultracentrifuge supernatant to isolate membrane fraction. Following solubilization membrane fraction with DDM detergent, centrifuge resuspend at 20,000×g for an hour at 4°C. Collect supernatant containing active LLOs. The crude membrane extract is active at 4°C for one week.

7. We use commercial N-terminal-TAMRA-GDQNATAF peptide substrate in our assay. In-house synthesized peptide with similar sequence can also be used as a glycosylation acceptor molecule.

8. The glycosylated protein will migrate slower in the SDS-PAGE gel due to the additional mass of the attached glycan, and on the anti-His immunoblot it will appear as a band slightly higher than the unmodified protein (if glycosylation efficiency is less than 100%, two bands will be apparent). The glycosylated form can be confirmed by appearance of a corresponding band on the glycan blot.

## 6. Acknowledgements

This work was supported by the Defense Threat Reduction Agency (GRANT11631647), the David and Lucile Packard Foundation, the Dreyfus Teacher-Scholar program, and the National Science Foundation (MCB 1413563). TJ was supported by a Royal Thai Government Fellowship. JCS was supported by the National Science Foundation Graduate Research Fellowship. JH was supported by an NDSEG Fellowship.

